# AI-Augmented R-Group Exploration in Medicinal Chemistry

**DOI:** 10.1101/2024.09.23.614417

**Authors:** Hongtao Zhao, Karolina Kwapień, Eva Nittinger, Christian Tyrchan, Magnus Nilsson, Susanne Berglund, Werngard Czechtizky

## Abstract

Efficient R-group exploration in the vast chemical space, enabled by increasingly available building blocks or generative AI, remains an open challenge. Here, we developed an enhanced Free-Wilson QSAR model embedding R-groups by atom-centric pharmacophoric features. Regioisomers of R-groups can be distinguished by explicitly accounting for the atomic positions. Good predictivity is observed consistently across 12 public datasets. Integrated into an open-source program, we showcase its application in performing classic Free-Wilson analysis as well as R-group exploration in uncharted chemical space.

---

R-group exploration is a key concept in medicinal chemistry that involves the systematic investigation of substituent groups (R-groups) on a molecular scaffold to optimize the pharmacological properties of a compound. This process is integral to the drug discovery and development pipeline, particularly during the hit-to-lead and lead optimization phases where chemists aim to improve the potency, selectivity, or other properties of a lead compound. Especially in the early phase, design of focused libraries or application of generative AI is chosen for resource-effective exploitation. It remains challenging to predict the exact impact of an R-group modification on the biological activity and pharmacokinetics. Quantitative structure-activity relationships (QSAR) modeling represents a powerful tool in this regard, employing molecular descriptors that encapsulate the spatial arrangement of R-groups to predict structure-activity relationships.^1-3^ However, the effectiveness of QSAR models is contingent upon their ability to accurately interpret the nuanced spatial arrangement of R-groups in relation to the heterogeneity of receptor binding sites. This necessitates extensive datasets to deconvolute such spatial complexities and generate reliable predictions.

Given the exponential increase in the available building blocks for library design,^4, 5^ it is in urgent need for a predictive model to help navigate the ultra-large chemical space enabled by R-group combinations. Building on the established R-group analysis, which decomposes a congeneric chemical series into a mapped core and associated R-groups, we have developed an AI-augmented R-group exploration program called AIR. Briefly, each atom within an R-group is encoded with a set of pharmacophoric and topological features, and subsequently arranged based on its topological distance (i.e., number of chemical bonds) to the anchoring point on the core (**Figure 1**). The embeddings of R-groups serve as the input for a feed-forward neural network. As a result, the spatial details of R-groups have been explicitly accounted for in predicting pharmacological properties. Our method has the ability to differentiate between regioisomers of R-groups, which sets it apart from the earlier Free-Wilson local QSAR model.^6^ This capability enables a more detailed exploration of chemical space, offering finer granularity in our analysis.

**Figure 1.**
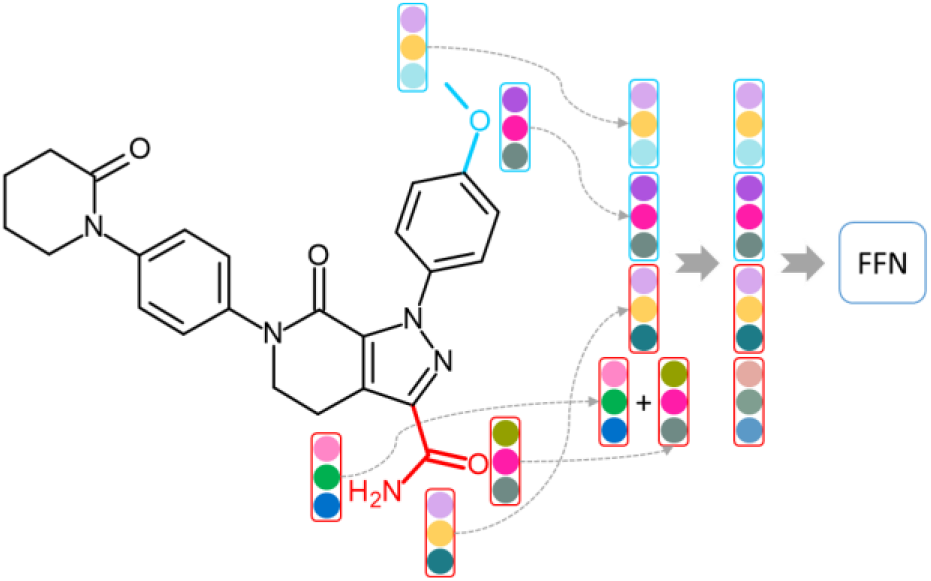
Schematics of the R-group based QSAR model. The core structure of the drug Apixaban is colored in black, and the two R-groups (i.e., the methoxy and primary amide) are in blue and red, respectively. Each atom in an R-group is encoded with a set of pharmacophoric and topological features into a vector of a fixed length. The encoded atoms are concatenated according to their topological distances from the core, and if there is more than one atom at the same distance, the vectors are added up. The embedded R-groups are then fed into a feedforward network (FFN) for property predictions.

The proposed method was tested on the 11 previous datasets extracted from chemical patents, encompassing a range of protein families including kinases (CDK5, JAK1, MAPK14, PIK3CA, and TYK2), serine proteases (F7 and PRSS2), a metalloproteinase (MMP2), a lipase (MGLL), a cytokine receptor (IL4), and a G protein-coupled receptor (GNRHR).^6^ In addition, a series of small-molecule inhibitors of DRD2 were retrieved from the ChEMBL database. Their core structures and R-group designations were summarized in **Table 1**.

**Table 1.**
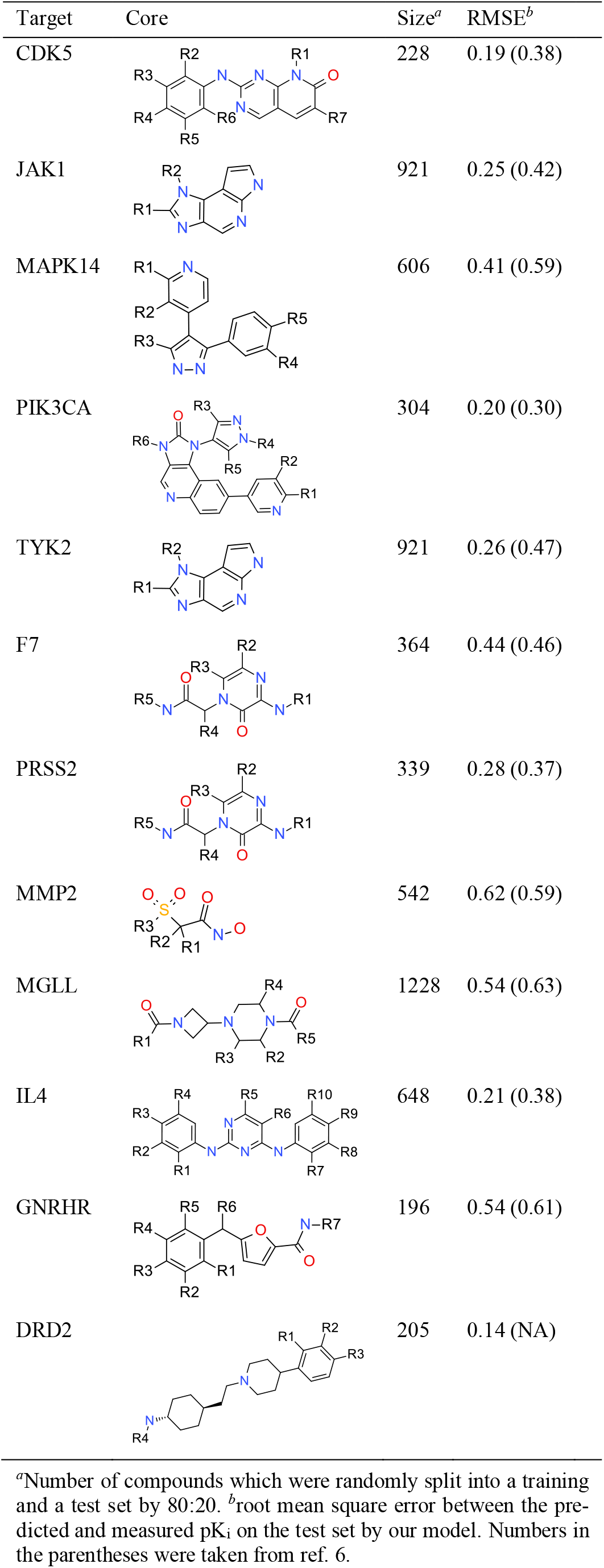
Model Performance on the 12 Datasets.

| Target | Core | Size<sup>a</sup> | RMSE<sup>b</sup> |
| --- | --- | --- | --- |
| CDK5 |  | 228 | 0.19 (0.38) |
| JAK1 |  | 921 | 0.25 (0.42) |
| MAPK14 |  | 606 | 0.41 (0.59) |
| PIK3CA |  | 304 | 0.20 (0.30) |
| TYK2 |  | 921 | 0.26 (0.47) |
| F7 |  | 364 | 0.44 (0.46) |
| PRSS2 |  | 339 | 0.28 (0.37) |
| MMP2 |  | 542 | 0.62 (0.59) |
| MGLL |  | 1228 | 0.54 (0.63) |
| IL4 |  | 648 | 0.21 (0.38) |
| GNRHR |  | 196 | 0.54 (0.61) |
| DRD2 |  | 205 | 0.14 (NA) |

<sup>a</sup>Number of compounds which were randomly split into a training and a test set by 80:20. <sup>b</sup>root mean square error between the predicted and measured $pK_i$ on the test set by our model. Numbers in the parentheses were taken from ref. 6.

First, we seek to evaluate its performance in predicting biological activities. Each dataset was randomly split into a training and a test set with a ratio of 4:1. Nine predictive models were trained on each training set by varying the random seeds. The mean of the ensemble predictions were plotted against the experimental values in **Figure 2**. The standard deviations of ensemble predictions provide a means to quantify the prediction uncertainties. The root-mean-square-error (RMSE) ranges from 0.14 to 0.62 with a median of 0.26 (**Table 1**). In comparison with the previous Free-Wilson local QSAR model,^6^ a significant improvement in the prediction accuracy was observed across nine datasets with similar performance on the remaining two (i.e., F7 and MMP2).

**Figure 2.**
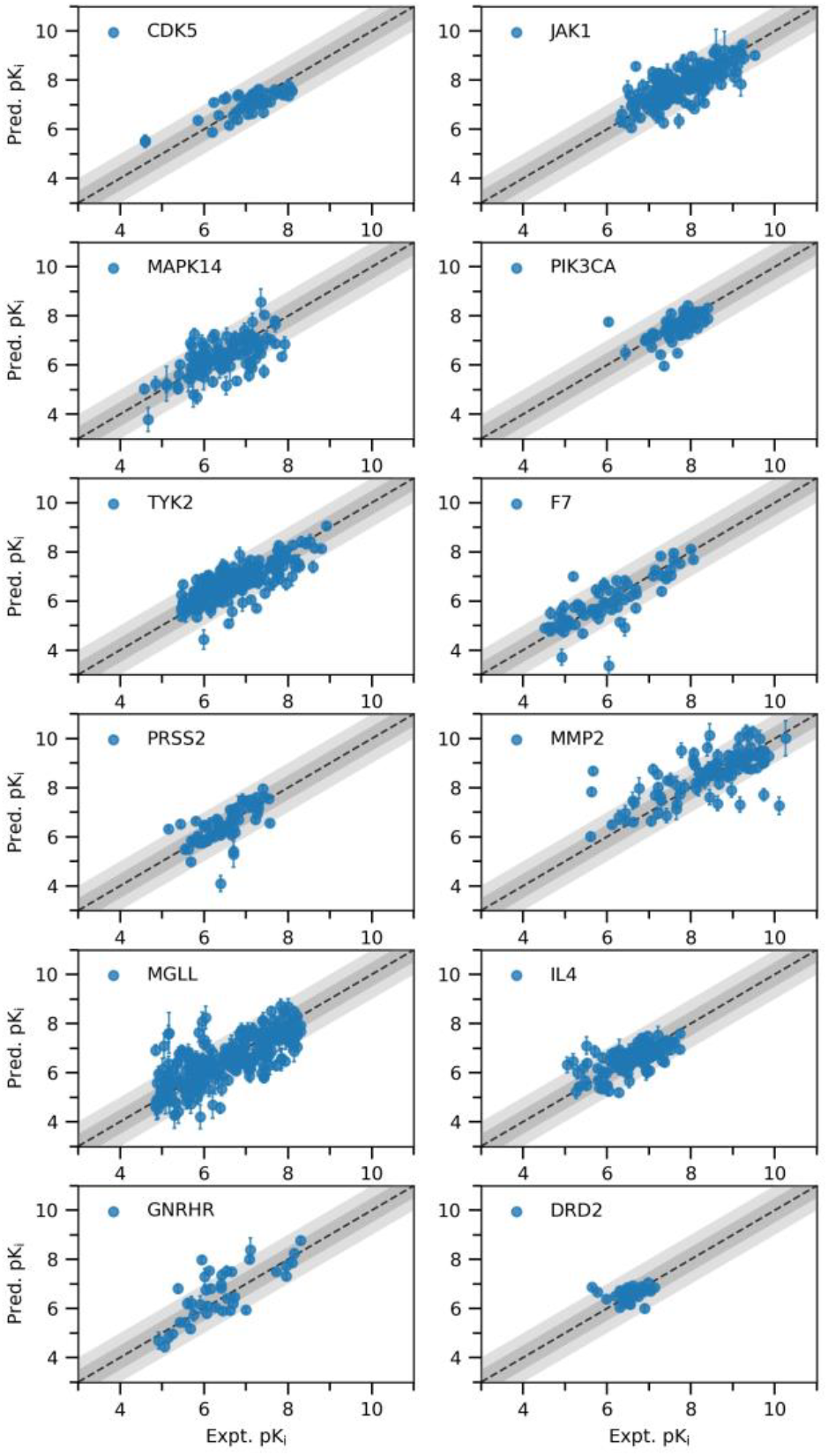
Scatter plots of the predicted against measured pK_i_ on the test sets from the 12 datasets, each of which was split into a training and a test set by 80:20. The error bars indicate the standard deviations of ensemble predictions by nine models.

To gain insight into prediction performance, we utilize t-SNE, a method that maps high-dimensional complex data into low-dimensional space while preserving the pairwise similarities. This allows us to understand the underlying patterns and relationships in the R-group embeddings in contrast to the full-molecule ECFP4 circular fingerprints (**Figure 3**). The ECFP4 fingerprints appear to be evenly distributed in space with no distinct separation between less active and more active compounds. In contrast, the R-group embeddings form clusters, wherein compounds with similar bioactivity tend to aggregate in space. This analysis suggests that the R-group embeddings may offer a more meaningful association with bioactivity compared to the ECFP4 fingerprints.

**Figure 3.**
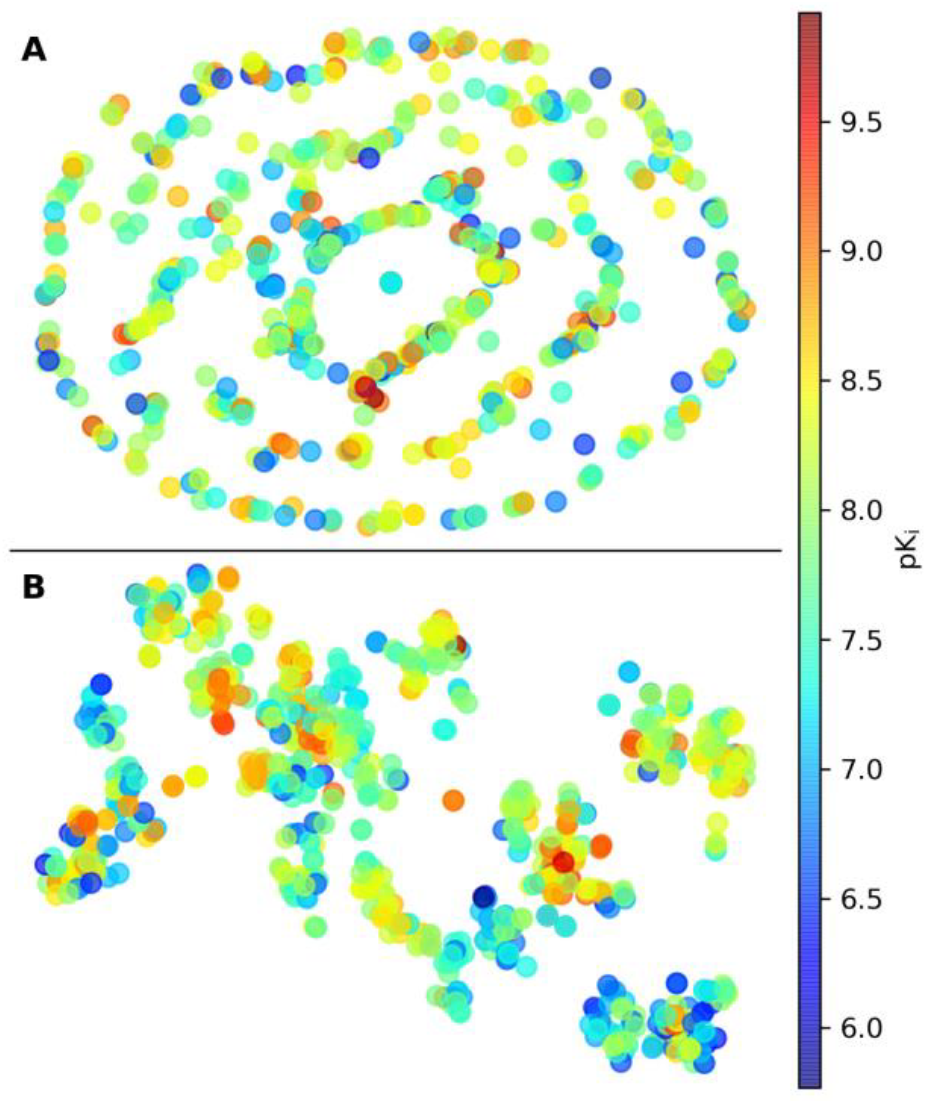
t-SNE visualization of the full-molecule ECFP4 fingerprints (**A**) and R-group embeddings (**B**) on the training set of JAK1 colored by pK_i_.

The method has been applied to several in-house projects over the past four years, demonstrating a positive impact on the overall progress. Due to the confidential nature of those early drug discovery projects, we opt to use the DRD2 dataset to illustrate its applications in the classic Free-Wilson analysis as well as exploration of large R-group space.

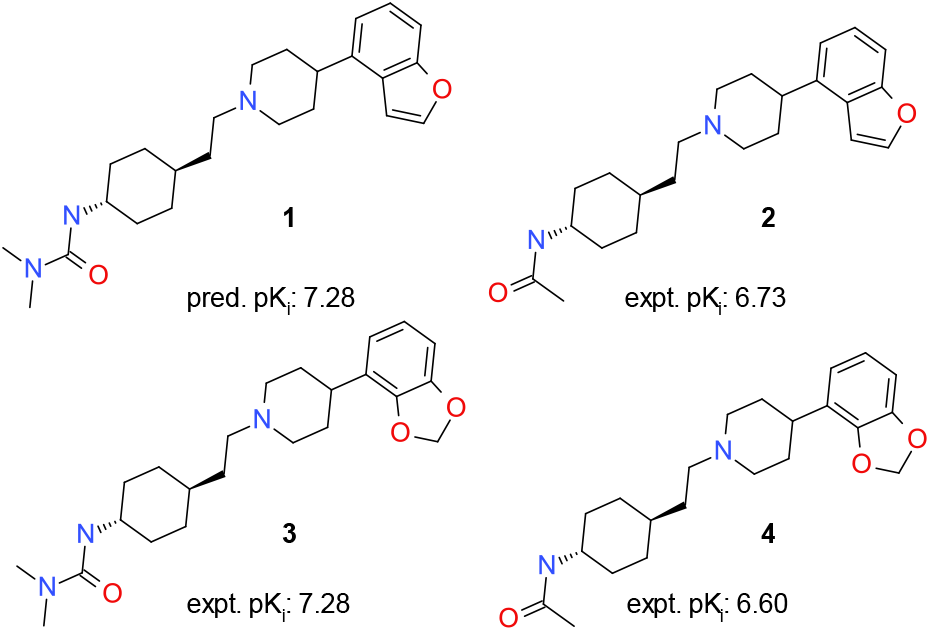

Free-Wilson analysis is a common practice in medicinal chemistry with the aim to understand the impact of individual R-groups on the overall activity of a molecule.^7^ Assuming that the contributions to activity made by R-groups at different substitution positions are additive, it enables the search for promising combinations of R-groups following a SAR exploration. We applied our method to the DRD2 series by performing an exhaustive combinatorial enumeration followed by activity prediction. Note that the program outputs actual chemical structures for the enumerated combinations, which is a desired feature for medicinal chemists. For instance, compound **1**, which is not present in the training set, was predicted to be equipotent as the most active compound **3**. These two compounds differ at the R^1^/R^2^ positions (refer to the core structure in **Table 1**). The matched molecular pair between compounds **2** and **4** suggests that the two substituents are equally potent, thus providing evidence for the predicted activity against **1**.

In Free-Wilson analysis, R-groups are treated as discrete and orthogonal to each other. Thus, it is impossible to predict the effect of a novel R-group not present in the training set even though it may represent a bioisostere to those already examined. With the pharmacophoric and topological embedding of R-groups, our method is capable of performing R-group exploration in uncharted chemical space. We applied a virtual amide coupling reaction to explore 15 522 R^4^ groups derived from the Enamine carboxylic reagents (**Figure 4**). Among the top-scored compounds, **5** and **6** were predicted to have a pK_i_ around 7.5. We turn to the matched molecular pair analysis^8-10^ for rationality behind the predictions. Compound **5** has a virtually identical molecular shape as compound **7**, and these two compounds differ in the substitution from 1,2,4-triazole to pyrrole. Note that 1,2,4-triazole has two more hydrogen bond acceptors than pyrrole. The matched molecular pair between **8** and **9** implies that a hydrogen bond acceptor at the meta-position of 3-pyridyl could improve the potency by 0.7 log unit. Both pyridine and 1,2,4-triazole have similar hydrogen bond basicity scale.^11, 12^ Taken together, compound **5** represents an interesting molecule for make in the next design cycle to explore the effect of a hydrogen bond acceptor at the meta-position being transferable or not. Compound **6** also shows a molecular shape similar to **10**, and both compounds have a hydrogen bond acceptor roughly in the same space. Note that the oxygen in isoxazole is a rather week hydrogen bond acceptor.^11, 12^ The predicted potency boost for **6** over **10** might be driven by the high lipophilicity of the CF_3_ group.

**Figure 4.**
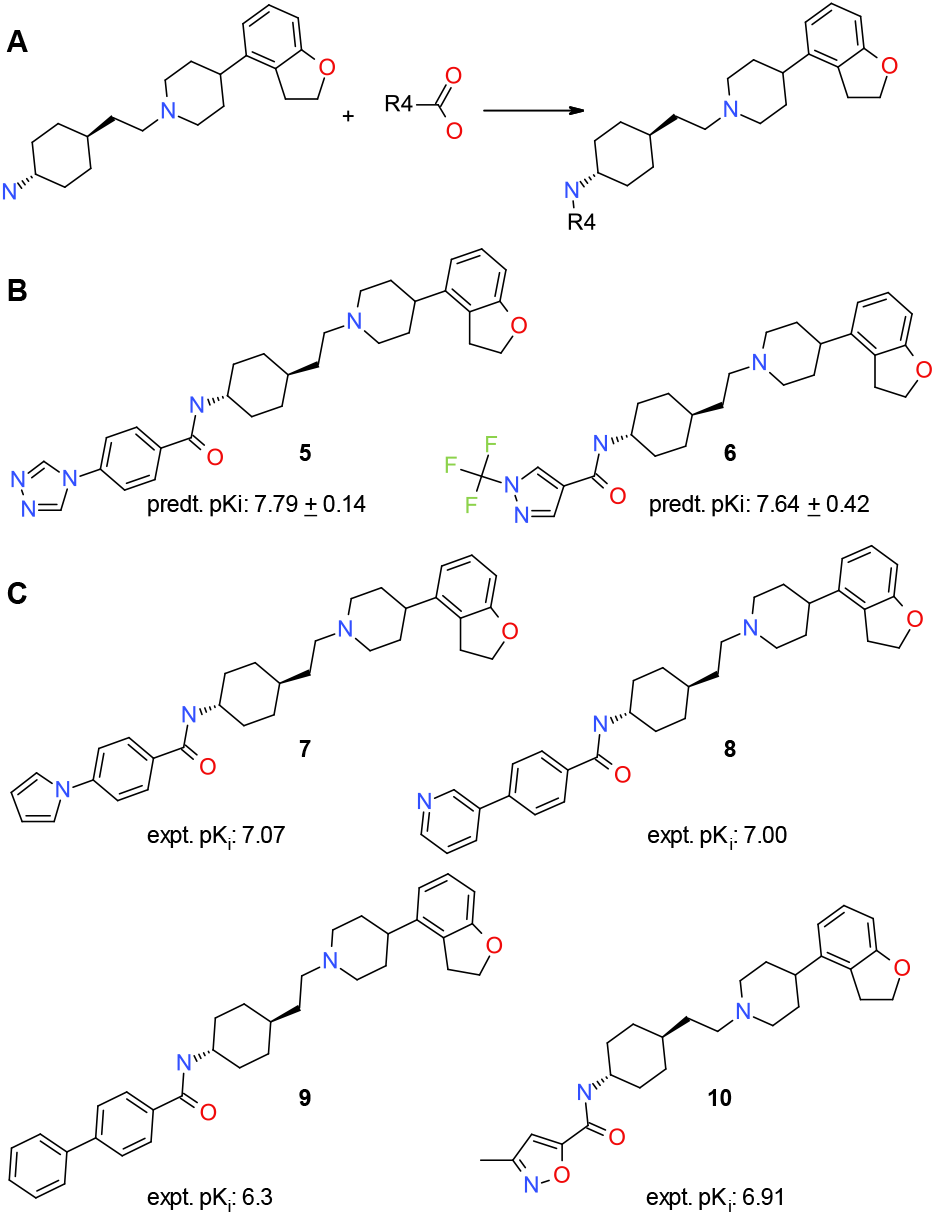
AI-augmented R-group exploration. (**A**) the reaction scheme of amide coupling, (**B**) two new compounds from the virtual library exploration, and (**C**) their nearest neighbors from those made.

The method was implemented using the R-group decomposition feature within the open-source RDKit project, which decomposes compounds into R-groups according to a user-defined core in a SMARTS string. A series of atom-centered pharmacophoric features, including characteristics such as hydrophobicity, ionizability, H-bond donor and acceptor properties, and chirality, were devised. Additionally, a set of patterns were engineered in relation to substitutions on a ring. Following the R-group decomposition, each atom in a designated R-group is encoded into a binary vector via substructure match with the pharmacophoric features and pattens (**Figure 5**). Furthermore, a radial atomic environment vector is generated for each heteroatom *i* of nitrogen or oxygen by

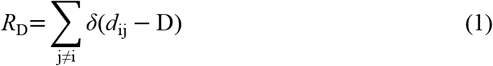

wherein *d*_ij_ is the topological distance between the heteroatom *i* and another atom *j* within the same R-group, *δ* is the delta function, and D∈{2, 3}. Then, an R-group at the *i*^th^ position (**r**_i_) is embedded by

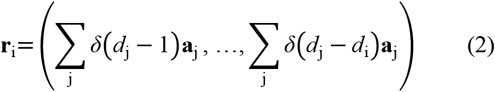

wherein *d*_j_ is the topological distance of the *j*^th^ atom to the anchoring point, **a**_j_ is the encoding of the *j*^th^ atom, *d*_i_ is the longest topological distance at the *i*^th^ position in the training set, and *j* runs over all atoms within the R-group. Finally, we generate property predictions *ý* by

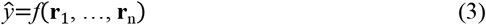

wherein *f*(·) is a machine-learning function and *n* is the number of substituting positions on the core. In comparison with the XGBoost, the feedforward neural network yielded a slight better performance with a single hidden layer of 16 neurons followed by a ReLU activation function. By sharing our open-source code, we hope that it could inspire the development of enhanced R-group embedding methods.

**Figure 5.**
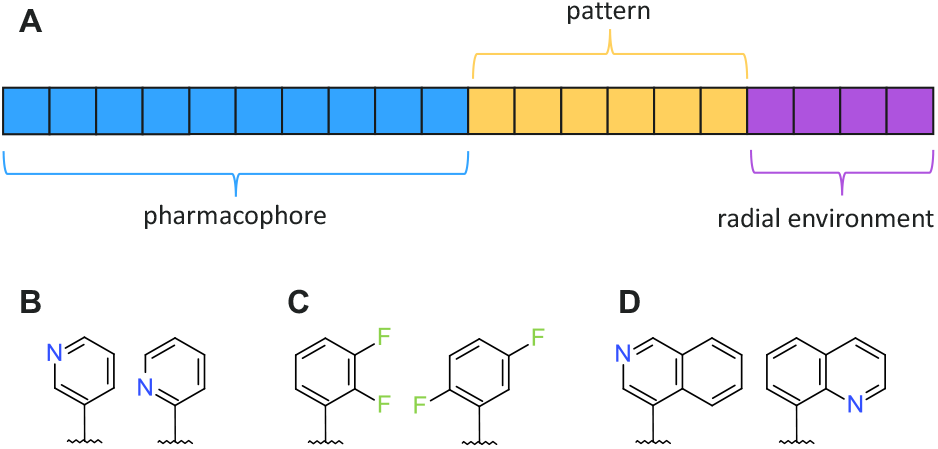
Atom encoding scheme. (**A**) atomic vector consisting of pharmacophores, patterns, and radial environment vectors of N and O, (**B**) example regioisomers distinguished by the topological distance of N to the anchoring point, (**C**) by ring substitution patterns (i.e., ortho-vs. para disubstitution), and (**D**) by radial environment vectors of N.

In summary, we have developed an improved Free-Wilson QSAR model that incorporates explicit encoding of R-groups along with atomic positions, and observed high prediction accuracy consistently across all 12 datasets. Implemented into the open-source program AIR, we showcase that it can perform the classic Free-Wilson analysis in search for promising combinations of already-explored R-groups, and that it can assist in the exploration of R-groups in uncharted chemical space.

## ASSOCIATED CONTENT

### Data Availability Statement

The source code of the python implementation and the dataset is available at https://github.com/WIMNZhao/AIR.

## AUTHOR INFORMATION

### Notes

H.Z., K.K., E.N., C.T., N.M., S.B. and W.C. are employees of AstraZeneca, and may own stock or stock options in AstraZeneca.

#### ABBREVIATIONS

QSAR,: quantitative structure-activity relationship;
AIR,: AI-augmented R-group exploration;
RMSE,: root mean square error.

---

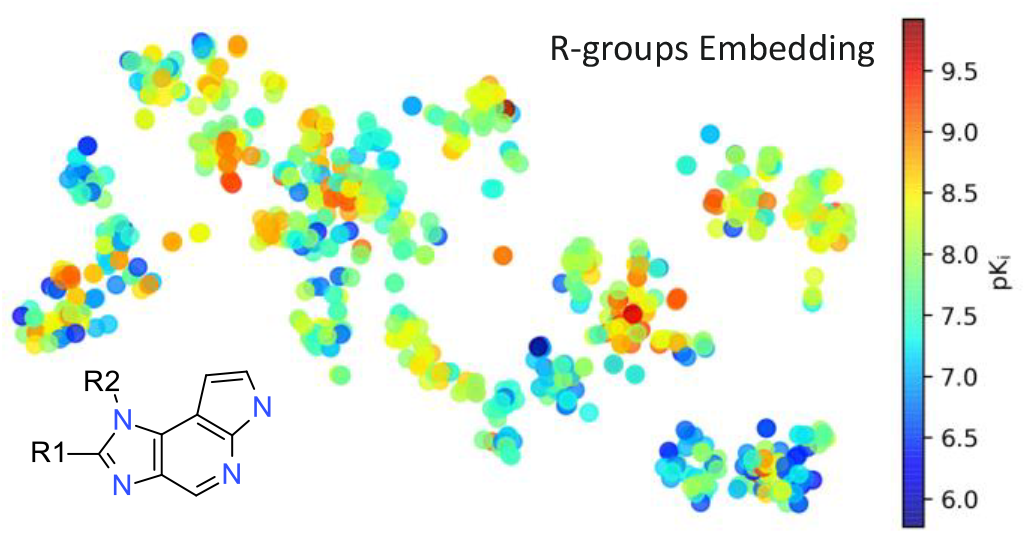
Insert Table of Contents artwork here

